# Medulloblastoma-associated mutations in the RNA helicase *DDX3X/DED1* cause defects in the translational response to TORC1 inhibition

**DOI:** 10.64898/2026.02.25.708058

**Authors:** Akanksha Swarup, Ryan A. Kuhs, Vanessa U. Hardman, Kennedy L. Howard, Shradha Subbaraman, Timothy A. Bolger

## Abstract

Medulloblastoma is the most common pediatric brain cancer, but current treatments are largely non-specific, often causing developmental side effects. Genomic sequencing identified the RNA helicase *DDX3X* as one of the most frequently mutated genes in this cancer and a potential treatment target, yet its role in tumor progression remains elusive. Prior studies have indicated that the mutations cause specific defects in translation; however, both DDX3X and its yeast ortholog Ded1 have also been associated with cellular stress responses, suggesting that the contribution of the *DDX3X* mutations to medulloblastoma might result from defects in the translational response to stress. Building on our prior study that replicated the *DDX3X* mutations in yeast *DED1* (*ded1-mam*), we examined the mutants’ effects following TOR pathway inactivation. First, we demonstrated that *ded1-mam* displayed substantial rapamycin-resistant growth compared to wild-type cells. In addition, similar to other *ded1* mutants, the *ded1-mam* had decreased degradation of Ded1 and the translation factor eIF4G1 under TOR inactivation. Notably, these differences did not result in increased bulk translation following rapamycin; rather, the growth phenotypes appeared to be driven by translation of specific mRNAs. Reporter assays demonstrated enhanced translation of mRNAs with unstructured 5′ UTRs in *ded1-mam* following TOR inhibition and a decrease in structured reporters. Furthermore, known Ded1 target genes with relatively unstructured 5’ UTRs showed upregulated protein levels in rapamycin. We thus hypothesize that mutant DDX3X selectively upregulates translation of unstructured, pro-growth transcripts while downregulating other structured transcripts, allowing tumor cells to bypass stress-induced growth controls and promoting medulloblastoma progression.

## Introduction

Medulloblastoma is the most common pediatric brain cancer, with around 400 new cases diagnosed in the United States each year (1, 2). Transcription profiling studies have classified medulloblastoma into four subtypes based on the affected signaling pathway: Wingless (WNT), Sonic Hedgehog (SHH), Group 3, and Group 4 (3). Current treatment strategies involve chemotherapy and radiation, and these procedures often result in long-term cognitive impairment and endocrine side effects (4). Therefore, more specific treatment strategies could help reduce the severe developmental effects of current broad-based treatments.

In recent years, deep sequencing studies have shown that the RNA helicase *DDX3X* is one of the most frequently mutated genes in medulloblastoma. These studies identified 42 different mutation sites in *DDX3X* in medulloblastoma patients, primarily associated with the WNT and SHH subtypes, and all of these mutations are single amino acid substitutions or deletions (Figure 1A) (4–8). DDX3X and its yeast ortholog Ded1 are members of the DEAD (Asp-Glu-Ala-Asp)-box protein family, which itself is part of the Superfamily 2 (SF2) of ATP-dependent RNA helicase proteins that are involved in ribonucleoprotein (RNP) remodeling (9, 10). Proteins that belong to the DEAD-box helicase family act in various roles in gene expression and RNA biology by unwinding double-stranded RNA and regulating RNP composition (9, 11, 12). The domain structure of DEAD-box proteins consists of a conserved helicase core region flanked by N- and C-terminal extensions that mediate regulatory and protein-protein interactions (Figure 1A)(11).

**Figure 1:**
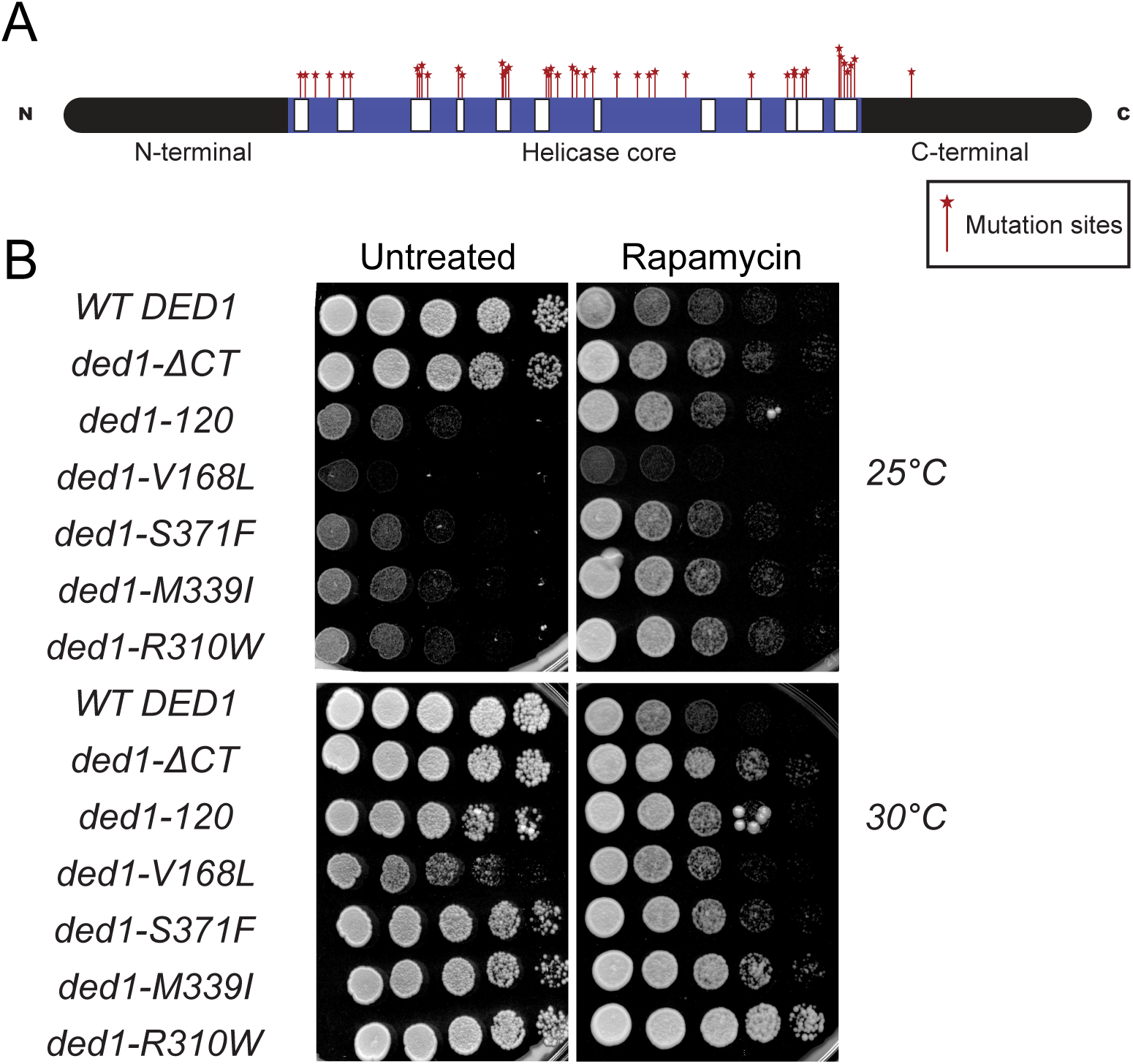
Medulloblastoma-associated mutations in *DDX3X/DED1* show rapamycin-resistant growth. (A) Domain schematic of Ded1/DDX3X with medulloblastoma-associated mutation sites (red lines) and conserved DEAD-box helicase motifs (white boxes). (B) Wild-type *DED1* cells and the indicated *ded1-mam* mutants were serially diluted onto nutrient-rich agar <u>in the presence or absence of</u> rapamycin (200 ng/ml) and incubated at 25 and 30°C.

DDX3X and Ded1 have conserved roles in eukaryotic translation initiation (10, 13). During initiation, translation factors and the 40S ribosomal subunit assemble on the mRNA to form the preinitiation complex (PIC) (14). DDX3X/Ded1 binds to components of the eukaryotic initiation factor 4F (eIF4F) complex, which recognizes the 5’ end of target mRNA during PIC assembly, and Ded1 is believed to help facilitate this process (15–17). Following assembly, the PIC will scan the mRNA in the 5’ to 3’ direction, in search of the start codon. In a large subset of mRNAs, secondary structures in the 5’UTR can impede this process, and DDX3X/Ded1’s most well-established role is to unwind these structures to facilitate start site scanning of these transcripts (18–22).

While DDX3X/Ded1’s role in translation initiation in nutrient-rich, pro-growth conditions is well-studied, its role in other conditions, such as cellular stress, is less understood. In stress, global translation is normally repressed by multiple pathways, including TOR, a central mediator of cell growth and division (23). Our prior work, primarily utilizing a *ded1* mutant that deletes its C-terminal region (*ded1-ΔCT*), suggested that when the TOR pathway is inactivated, Ded1 acts as a translation repressor, helping the cell to halt growth and conserve energy during stress (24). Furthermore, this repression was at least partly mediated by the removal of the critical scaffolding protein eIF4G1 from translation complexes by Ded1, followed by the degradation of both proteins (24). However, DDX3X/Ded1’s role in stress appears to be complex. For example, in heat shock, it has been reported that Ded1 is repressed and unable to translate housekeeping genes with complex 5’ UTRs (25). These studies and others have suggested a model where DDX3X/Ded1 decreases growth and bulk translation while increasing translation of stress mRNAs in the initial stress response, followed by a role in the resumption of growth during stress adaptation and recovery (26). Taken together, these studies show that DDX3X/Ded1 plays an important role in the cellular stress response. Notably, stress responses are implicated in oncogenesis; tumor cells are frequently under stress, and thus, cancer drivers include alterations that allow cells to overcome stress-mediated growth inhibition (27).

In line with its involvement in brain cancer, DDX3X is also essential for normal brain development. *Ddx3x* is expressed throughout the developing brain in mouse embryos, and tissue-specific knockouts result in impaired neural progenitor growth, decreased cortical generation, reduced neuronal differentiation, microcephaly, and apoptosis (28, 29). Furthermore, *Ddx3x* is critical for maintaining the balance between progenitors and neurons, regulating hindbrain patterning and development (30).

The 42 different mutations in *DDX3X* associated with medulloblastoma (see Figure 1A) are spread across its conserved helicase core (31). Several studies have characterized small subsets of these mutations, and they have generally been found to impair translation. Protein levels of a known Ded1 target were decreased in a handful of medulloblastoma-associated DDX3X mutants in *S. pombe* (32), and overexpression of several DDX3X medulloblastoma mutant strains have displayed bulk translation inhibition (33, 34). However, many of the mutants do not appear to have defects in general translation, implying that the mutations do not primarily affect global translation but rather affect translation of specific genes (31).

Despite this prior research, a knowledge gap persists regarding the role of *DDX3X* mutations in cancer progression and the molecular mechanism behind this process. In particular, few studies have taken a broad approach to characterizing the spectrum of different mutations identified in medulloblastoma. We recently characterized all 42 mutants by leveraging the genetic tractability of yeast to generate mutants that replicate each of the 42 identified *DDX3X* mutations in its yeast ortholog, *DED1* (31). These *ded1-medulloblastoma-associated-mutants* (*ded1-mam*) displayed various growth and translation phenotypes. Still, the most common phenotype was a defect in translation of reporter mRNAs containing structured 5’UTRs, suggesting that defective translation of this class of mRNAs may be responsible for the effect of the *DDX3X* mutations in medulloblastoma. However, this prior work did not consider the role of DDX3X/Ded1’s function in cellular stress. Here, we examined the potential effect of the medulloblastoma-associated mutations on DDX3X/Ded1 stress function by again utilizing our *ded1-mam* lines. As shown below, most of the mutants demonstrated a rapamycin-resistant growth phenotype relative to wild-type, similar to our prior work with *ded1-ΔCT.* However, the mutants still displayed bulk downregulation of translation under stress, suggesting that more specific translation defects are responsible. Working to investigate these phenotypes, we found that the mutants have increased persistence of Ded1 and eIF4G1 following TOR inactivation. Furthermore, we found that the medulloblastoma-associated mutants showed increased translation of reporter mRNAs with unstructured 5’ UTRs and decreased translation of mRNAs with structured 5’ ones upon TOR inactivation. Likewise, protein levels of selected pro-growth, relatively unstructured targets of Ded1 were also increased in the mutant cells under stress. These results point to a prospective role of DDX3X/Ded1 in preferential unwinding and translation of certain mRNAs depending on 5’UTR structure in combination with the availability of eIF4G1 to stimulate translation during stress conditions. We suggest that in medulloblastoma, *DDX3X*’s role in stress response is dysregulated by the identified mutations, allowing medulloblastoma cells to continue to proliferate in adverse conditions and contributing to tumor progression.

## Results

### Medulloblastoma-associated mutations in *DED1/DDX3X* improve growth following TOR inhibition

Prior studies have identified 42 different mutation sites in *DDX3X* associated with medulloblastoma progression, all of which are in the central helicase domains (Figure 1A) (4–8). As described previously, we have taken advantage of the high degree of sequence homology in this region to construct yeast strains with the corresponding mutations in *DED1*, terming these *ded1 <u>m</u>edulloblastoma-<u>a</u>ssociated <u>m</u>utants* (*ded1-mam*) (31). Out of these 42 mutants, 9 of them were lethal, while the remaining *ded1-mam* mutants had varied growth and translation phenotypes consistent with established functions of Ded1/DDX3X (31). However, other than a moderate increase in stress granules in some mutants, whether the functions of Ded1/DDX3X in stress responses are affected by these mutations is not yet clear (31, 34).

In order to begin to characterize the effects of these mutations during cellular stress, we screened the 33 viable *ded1-mam* mutants for changes in growth following TOR pathway inactivation via rapamycin, a specific inhibitor of TORC1 (35). Plated on nutrient-rich media in the absence of rapamycin, all 33 viable *ded1-mam* strains showed at least some growth at 30°C, and 16 displayed cold-sensitive growth defects at 25°C (Table 1), consistent with our previous study (31). Representative growth assays are shown in Figure 1B, and a complete set with all tested strains is shown in Supplementary Figures S1 and S2. Upon addition of rapamycin (200 ng/ml) to the media, growth of wild-type *DED1* cells was markedly inhibited at both 25°C and 30°C, as expected (Figure 1B). In contrast, two previously characterized *ded1* mutants, *ded1-*Δ*CT* and *ded1-120*, that delete the C-terminal region and impair Ded1 helicase activity, respectively, grew better than the wild-type *DED1* strain in the presence of rapamycin at both temperatures, as shown previously (13, 17, 24). Remarkably, we observed that most of the *ded1-mam* mutants (21 of 33) displayed a similar rapamycin-resistant phenotype (Table 1 and Supplementary Figures S1 & S2). Notably, many of the cold-sensitive growth phenotypes in the *ded1-mam* mutants were completely reversed in the presence of rapamycin (Figure 1B). There is also a clear correlation between the cold-sensitive growth defect in these strains in rich media and the extent of rapamycin-resistant growth (Supplementary Figure S3). The increased growth of *ded1-mam* compared to wild-type *DED1* strains in rapamycin suggests that these mutants have defects in responding to cellular stress, similar to the *ded1-*Δ*CT* and *ded1-120* strains. Furthermore, roughly two-thirds of *ded1-mam* mutants were at least partially rapamycin-resistant, suggesting that alterations in the function of Ded1/DDX3X in stress responses may be a common phenotype underlying the contributions of these mutations to medulloblastoma progression.

**Table 1:** Rapamycin-resistant phenotype and temperature sensitivity of DDX3X/DED1 medulloblastoma-associated mutations. The identified *DDX3X* mutations organized by amino acid sequence with the orthologous change in the conserved Ded1 sequence. Growth assays of the *ded1-mam* strains were performed at 25 and 30°C on rich media in the presence or absence of rapamycin (200 ng/ml). Growth phenotypes: no growth (-), slow growth (+), moderate growth (++), and growth equivalent to wild-type *DED1* yeast without rapamycin (+++). ND = not determined, mutation is lethal at all temperatures. Green = rapamycin-resistant *ded1-mam* strain.

| **DDX3X**<br>**mutation**<br>**site** | **DED1** | **Amino acid**<br>**substitution** | **Temperature-**<br>**sensitive**<br>**growth?** | **Rapamycin-**<br>**resistant at**<br>**25°C?** | **Rapamycin-**<br>**resistant at**<br>**30°C?** |
| --- | --- | --- | --- | --- | --- |
| *DDX3X* |  |  | +++ | - | - |
| - | *ded1-ΔCT* |  | +++ | + | + |
| - | *ded1-120* |  | - | +++ | ++ |
| 204 | 166 | T/A | ++ | + | - |
| 206 | 168 | V/L | + | ++ | +++ |
| 214 | V176 | I/S,KV | ND | ND | ND |
| 222 | 184 | A/P | +++ | - | - |
| 227 | 189 | G/V | +++ | - | - |
| 231 | 193 | T/A | + | +++ | ++ |
| 275 | 234 | T/M | ++ | + | - |
| 276 | 235 | R/K,T | + | +++ | +++ |
| 277 | 236 | E/Q | +++ | - | - |
| 281 | 240 | Q/H | ND | ND | ND |
| 302 | 261 | G/V | ND | ND | ND |
| 303 | 262 | G/V | + | +++ | +++ |
| 325 | 284 | G/E,R | ND | ND | ND |
| 326 | 285 | R/C | ++ | +++ | ++ |
| 327 | 286 | L/V | +++ | - | - |
| 329 | 288 | D/V | +++ | + | - |
| 351 | 310 | R/W | + | +++ | +++ |
| 353 | 312 | L/F | ++ | + | + |
| 354 | 313 | D/H,V | +++ | + | - |
| 357 | 316 | F/S | ++ | ++ | ++ |
| 368 | 326 | D/V | +++ | - | - |
| 370 | 329 | M/R | +++ | + | - |
| 376 | 335 | R/C,S | +++ | - | - |
| 380 | 339 | M/I | + | +++ | ++ |
| 384 | 343 | T/P | ND | ND | ND |
| 393 | 352 | A/S,V | +++ | - | - |
| 405 | 364 | V/L | +++ | - | - |
| 412 | 371 | S/F | + | +++ | ++ |
| 415 | 374 | I/- | ND | ND | ND |
| 434 | 393 | L/- | +++ | + | - |
| 475 | 433 | R/C | +++ | + | + |
| 496 | 454 | V/M | +++ | + | - |
| 501 | 459 | A/- | ND | ND | ND |
| 504 | 462 | G/V | +++ | ++ | ++ |
| 506 | 464 | D/Y | +++ | - | - |
| 527 | 485 | H/Y | ND | ND | ND |
| 528 | 486 | R/H | ++ | - | - |
| 530 | 488 | G/A | ++ | - | - |
| 532 | 490 | T/M | +++ | - | - |
| 534 | 492 | R/S,H,C | ND | ND | ND |
| 536 | 494 | G/R | ++ | + | - |
| 568 | 526 | P/L | +++ | - | + |

### Medulloblastoma-associated mutants of *DDX3X/DED1* display partially impaired translation downregulation following TOR inhibition

As discussed above, DDX3X/Ded1 has established roles in both stimulating and repressing translation in different conditions (Chuang et al., 1997; Lai et al., 2008; Aryanpur et al., 2019; Hilliker et al., 2011; Iserman et al., 2020; Shih et al., 2008; Aryanpur 2022). In particular, our prior research has shown that Ded1 downregulates bulk translation following TORC1 inhibition, and this downregulation is impaired in some *ded1* mutants (24). Therefore, we wanted to observe whether the *ded1-mam* mutants have defects in downregulating bulk translation in a similar fashion. To test this, we performed polyribosome profiling to determine the overall translation status of two *ded1-mam* mutants, *ded1-F316S* and *-R310W*, relative to wild-type *DED1* during rapamycin treatment. The mutant and wild-type cells were treated for 40 minutes with rapamycin, and profiles were generated by measuring the RNA concentration (Abs_254nm_) in cellular extracts subjected to sucrose density gradient centrifugation (Figure 2A-F). The polysome to 80S monosome (P/M) ratio was then calculated from the profile for each condition as a measure of translational activity (Figure 2G) (31, 39, 40).

**Figure 2:**
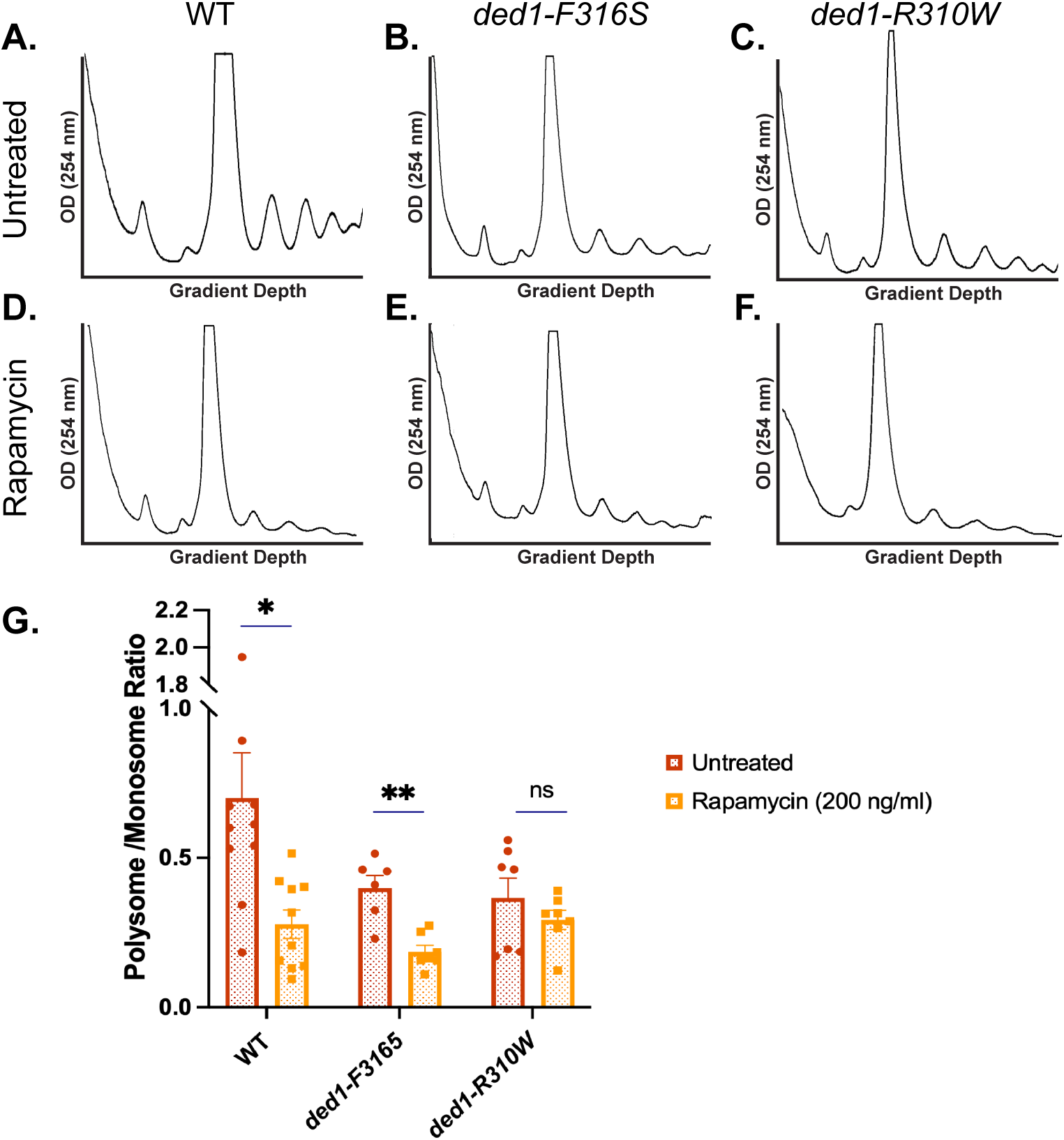
Medulloblastoma-associated mutants of DDX3X/DED1 downregulate translation following TOR inhibition. (A-F) Polyribosomal profiles of wild-type and the indicated ded1-mam mutants grown in the presence of rapamycin for 40 minutes, following sucrose density centrifugation and OD254nm analysis of cell lysates. (G) The polysome/monosome (P/M) ratios were determined by comparing the sum of the areas of the polysome peaks with the area of the monosome peak. For each strain/condition, the mean P/M ratio, the SEM, and the individual replicates is shown. * p < 0.05, ** p < 0.01, ns = not significant vs. the untreated condition for each strain.

The polyribosome profile of untreated wild-type cells showed robust translation activity, as expected (Figure 2A). However, translation was somewhat reduced in both *ded1-mam* mutants in this condition with P/M ratios that were lower than that of wild-type *DED1* (Figure 2B, C & G). This result is consistent with our prior study in which both *ded1-F316S* and *-R310W* showed moderately reduced P/M ratios after a shift to non-permissive temperatures, suggesting a partial defect in bulk translation (31). After rapamycin treatment, bulk translation decreased substantially in wild-type *DED1* cells, as reflected in both the profiles and the P/M ratio, which decreased from 0.69 to 0.26 (Figure 2D & G), reflecting the general repression of translation in these conditions (41). Likewise, the bulk translational activity of both *ded1-F316S* and *-R310W* also declined in rapamycin (Figure 2E & F). Interestingly, however, the magnitude of the reductions in the mutants is not as great as in wild-type cells, resulting in P/M ratios (0.19 and 0.29) that were similar to the rapamycin-treated wild-type cells (Figure 2G). This reduction in the degree of translation inhibition in the *ded1-mam* mutants following TORC1 inhibition is similar to that observed previously in the *ded1-ΔCT* mutant (24). That being said, since there is little difference in the raw P/M ratio between the mutants and wild-type in these conditions, it is unlikely that the rapamycin-resistant growth phenotype observed in the *ded1-mam* mutants is due to a general increase in bulk translation. Rather, more specific changes in translation regulation may be responsible for this stress-resistant growth.

### Medulloblastoma-associated mutants of *DDX3X/DED1* demonstrate reduced degradation of Ded1 and eIF4G1 upon TOR inactivation

Under normal conditions, it has been observed that Ded1 promotes 48S PIC formation and interacts with eIF4G1 via its C-terminus to support translation and cell growth (16, 17, 36). However, we had previously proposed a model where, under stress, wild-type Ded1 dissociates eIF4G1 from the PIC, leading to co-degradation of both proteins and translation repression (Aryanpur et al., 2019). In the *ded1-ΔCT* mutant, this remodeling of the PIC is reduced, and Ded1 and eIF4G1 degradation is slowed, resulting in increased translation and rapamycin-resistant growth. We predicted that a similar reduction in degradation would be observed in the *ded1-mam* mutants, which might contribute to their increased growth under stress conditions. To demonstrate this, we treated wild-type *DED1*, *ded1-*Δ*CT*, and six of the *ded1-mam* mutants (*ded1-V168L*, *-R235K*, *-G262V*, *-R310W*, *-F316S,* and *-M339I*) with rapamycin for 4 hours. Cell extracts were prepared and run on SDS-PAGE gels to analyze levels of Ded1 and eIF4G1 in untreated and rapamycin-treated cells (Figure 3A). In wild-type *DED1* cells, we observed that Ded1 and eIF4G1 levels were higher in untreated conditions compared to after rapamycin treatment (Figure 3A). Following rapamycin treatment, a sharp decrease in the levels of both Ded1 and eIF4G1 was seen, as expected (Figure 3B & D) (24). By contrast, in the *ded1-*Δ*CT* mutant, while there was still a decrease in Ded1 and eIF4G1 levels after rapamycin treatment, the protein levels were higher than in treated wild-type cells (Figure 3A, B, & D), consistent with our prior studies (24).

**Figure 3:**
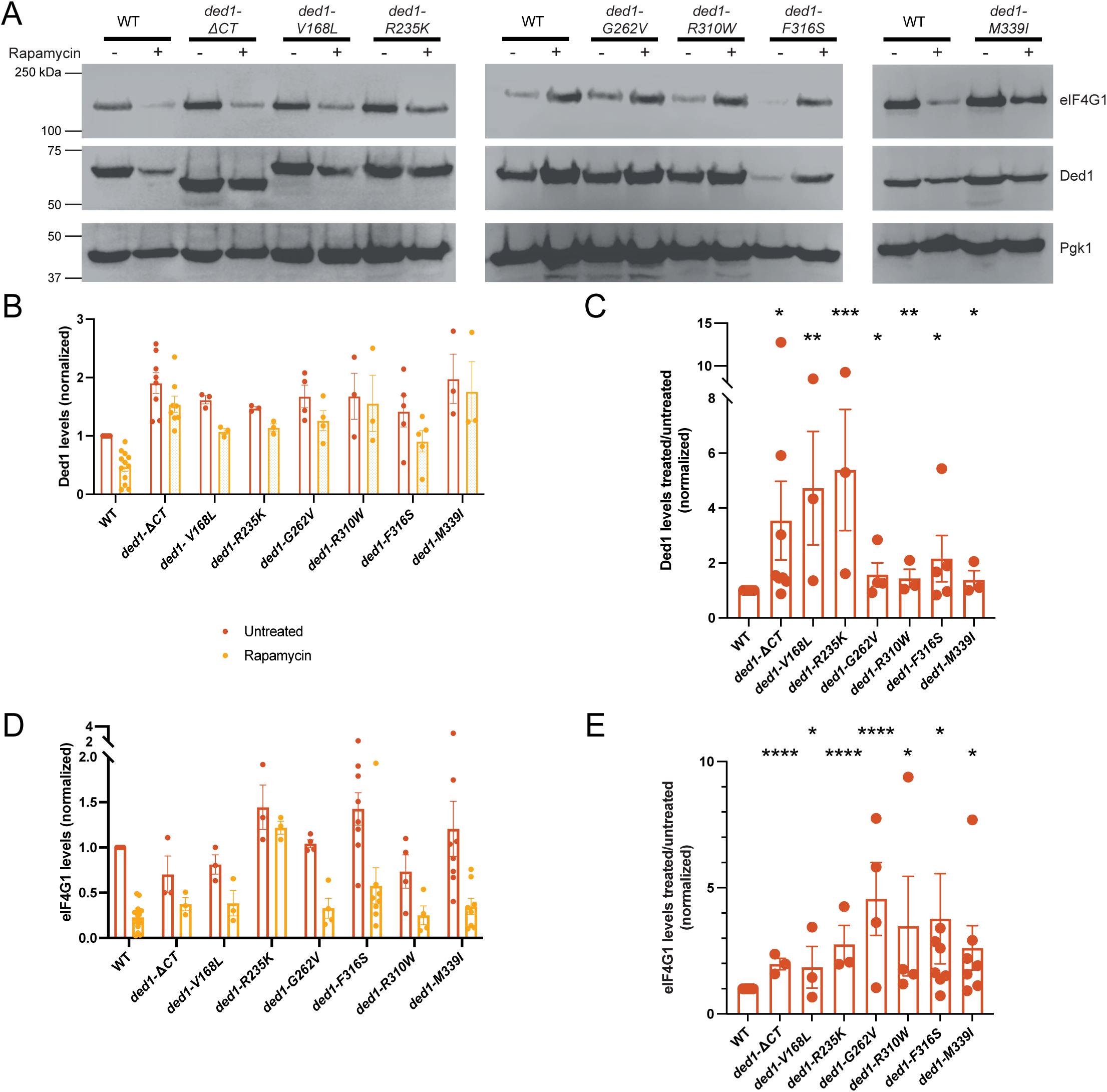
Medulloblastoma-associated mutants of *DDX3X/DED1* show reduced degradation of Ded1 and eIF4G1 upon TOR inactivation. (A) Protein extracts from wild-type *DED1* cells and the indicated *ded1-mam* mutants were treated with rapamycin (200 ng/ml) or not, then cell extracts were run on SDS-PAGE and blotted using anti-Ded1, anti-eIF4G1 (Tif4631), and anti-Pgk1 (loading control). Representative images are shown. (B, D) Band densitometry was performed, and Ded1 and eIF4G1 protein levels, respectively, are shown, normalized to levels in untreated wild-type *DED1* cells. (C, E) Fold change in levels of Ded1 and eIF4G1, respectively, are shown following rapamycin treatment compared to untreated samples. Values are normalized to the fold change in wild-type *DED1* cells. *\* p* < 0.05, *\*\* p* < 0.01, \**\*\* p* < 0.001, *\*\*\*\* p* < 0.0001 vs. wild-type *DED1*. For (B-E), individual replicates, means, and SEM are shown.

For the *ded1-mam* mutants, we observed phenotypes similar to *ded1-ΔCT*, wherein each of the mutants showed decreased degradation (that is, increased levels) of Ded1 and eIF4G1 compared to wild-type cells following rapamycin treatment (Figure 3A, B & D). In several mutants, the fold change in Ded1 levels was less than in *ded1-ΔCT*, though still significant (Figure 3C). For eIF4G1, however, the fold change in protein levels after treatment in the *ded1-mam* mutants was similar to or greater than that observed in *ded1-ΔCT* cells (Figure 3E). These differences may be partly explained by moderate changes in basal Ded1 and eIF4G1 levels in untreated cells, resulting in a different fold change calculation, even though higher levels of protein clearly remain post-treatment. Consistent with prior studies, only minor changes in the protein levels (<2-fold) were observed in untreated cells (31, 34). Thus, Ded1 and eIF4G1 show decreased degradation upon rapamycin treatment in the *ded1* medulloblastoma-associated mutants, similar to our previous observations of *ded1-ΔCT* (24). These results suggest that the increased amount of Ded1 and eIF4G1 is responsible for the increase in rapamycin-resistant growth in these mutants and may cause altered translation regulation in these conditions.

### Medulloblastoma-associated mutants of *DDX3X/DED1* show opposing effects on translation in stress conditions depending on 5’UTR structure

Both DDX3X and Ded1 have documented roles in facilitating start site scanning via unwinding secondary structures in 5’ UTRs of mRNAs (Guenther et al., 2018; Lai et al., 2008; Sen et al., 2015; Soto-Rifo et al., 2012). To study how this process is affected in *ded1-mam,* we obtained previously published luciferase reporters with 5’ UTRs derived from *RPL41A* that either contain a stem loop (ΔG_free_ = −3.7 kcal/mol) or are unstructured (Sen et al., 2015) (Figure 4A). In our previous study, we examined the activity of these reporters in a subset of *ded1-mam* mutants in normal conditions, where the mutants showed consistently reduced translation of the structured reporter compared to the unstructured one, implying that the *ded1-mam* mutants were defective in unwinding of RNA structures in the 5’ UTR (31). These specific translation defects also correlated with the mutants’ cold-sensitive growth defects. How the *ded1-mam* mutants behave under stress conditions is unknown, however.

**Figure 4:**
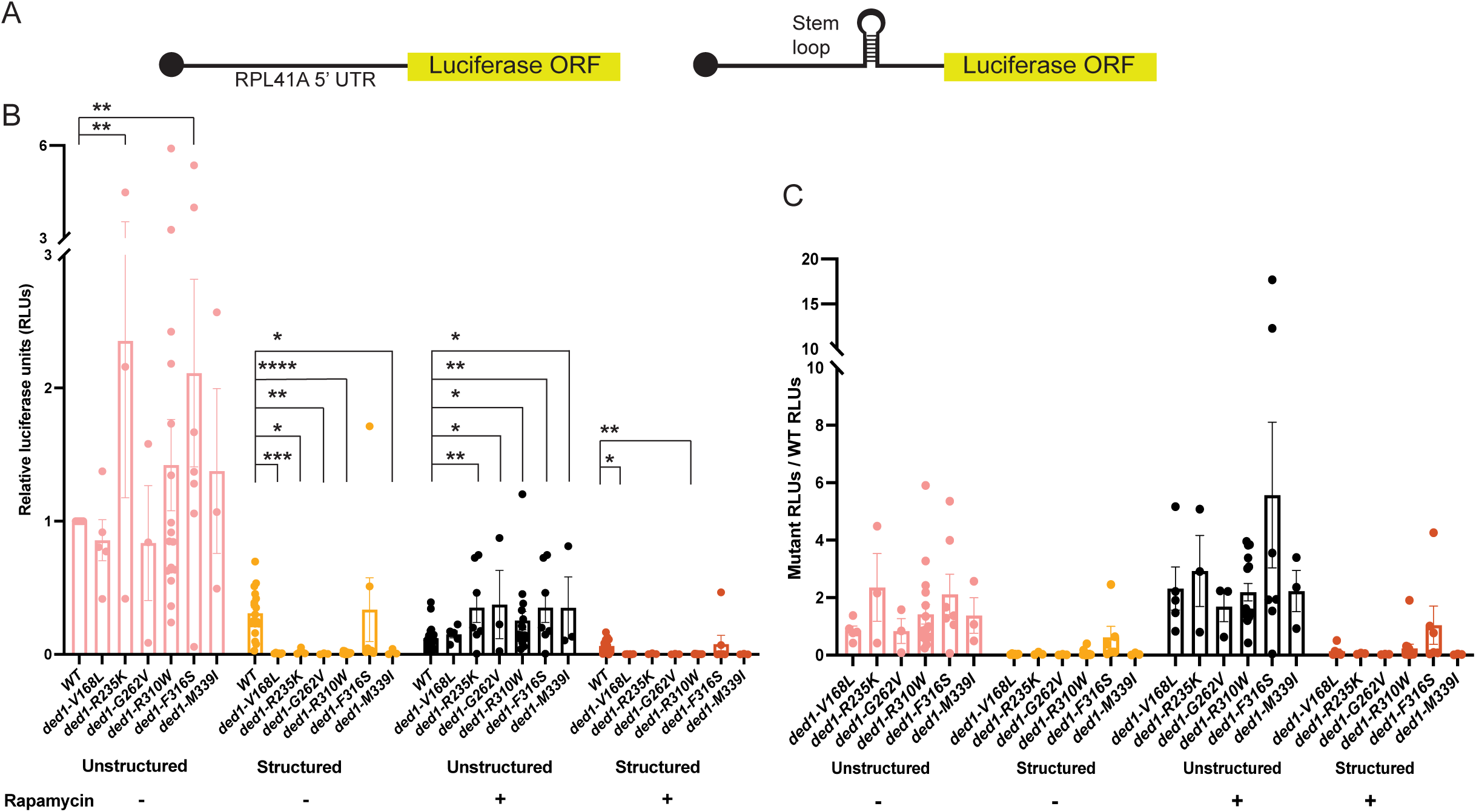
Medulloblastoma-associated mutants of *DDX3X/DED1* show opposing effects on translation in stress conditions depending on 5’UTR structure. (A) Schematic of the unstructured (left) and structured (right) 5’UTR luciferase reporters. The 5’UTRs are derived from that of RPL41A, lengthened to 91 nt, with a 3.7 kJ/mol stem loop inserted in the structured reporter. (B) Wild-type *DED1* cells and the indicated *ded1-mam* mutants containing these reporters were treated with rapamycin (200 ng/ml) or not for 4 hours, and cell extracts were subjected to luciferase assays. The relative luciferase units (RLUs) are shown, normalized to untreated wild-type cells. (C) Fold change in RLUs, relative to the values for wild-type cells, for each reporter in each *ded1-mam* mutant treated with rapamycin or not. *\* p* < 0.05, *\*\* p* < 0.01, *** *p* < 0.001, **** *p* < 0.0001 vs. wild-type *DED1* with the same reporter in the same condition.

Based on the growth phenotypes, changes in Ded1 and eIF4G1 levels, and the lack of a substantial phenotype in bulk translation in the *ded1-mam* mutants during TOR inactivation, we hypothesized that alterations in scanning through structured and unstructured 5’UTRs may be responsible for the increased growth in these conditions. To address this hypothesis, we transformed the structured and unstructured reporters into wild-type *DED1* and the indicated *ded1-mam* mutants, treated the cells with rapamycin for 4 hours, and conducted luciferase assays (Figure 4B). As expected, we observed that in wild-type *DED1* cells, luciferase activity from the structured reporter was lower than that from the unstructured reporter in both untreated and rapamycin-treated conditions (Brown et al., 2021; Sen et al., 2015). Furthermore, as previously reported, the *ded1-mam* showed significantly reduced luciferase activity in structured reporters compared to wild-type *DED1* under untreated conditions (Figure 4B & C)(31). For the unstructured reporters in untreated conditions, the activity in *ded1-mam* cells was generally similar to wild-type *DED1*, although it was somewhat increased in *ded1-R235K* and *ded1-F316S* (Figures 4B & C).

The effect of rapamycin, however, has not been tested previously with these reporters. As expected, based on the general repression of translation following rapamycin treatment (see Figure 2) (43), both unstructured and structured reporter activity was decreased upon rapamycin treatment in wild-type *DED1* cells (Figure 4B). Consistent with the effect shown in untreated conditions, the activity from the structured reporters in *ded1-mam* cells also seemed to decrease compared to wild-type *DED1* under rapamycin treatment, although due to the very low activity, this difference was insignificant for some mutants (Figure 4B & C). Strikingly, however, unstructured reporter activity was significantly higher in the *ded1-mam* mutants compared to wild-type *DED1* cells after rapamycin treatment (Figure 4B), with increases from approximately 2 to 5-fold across the mutants (Figure 4C). This result is intriguing since it implies a sharp contrast in the way that different classes of mRNAs are translated in *ded1-mam* mutants compared to wild-type cells. Specifically, mRNAs with unstructured 5’ UTRs are significantly *upregulated* in the *ded1-mam* mutants relative to wild-type *DED1* under stress, and mRNAs with structured 5’UTRs are *downregulated* in the *ded1-mam* cells. Moreover, this data suggests that the rapamycin-resistant growth of *ded1-mam* cells may be driven by the translation of mRNAs with relatively unstructured 5’ UTRs, while mRNAs with relatively structured 5’ UTRs may be downregulated.

### Medulloblastoma-associated mutants of *DDX3X/DED1* have increased protein levels of known *DED1* targets with unstructured 5’ UTRs following TOR inactivation

The luciferase assays shown in Figure 4 demonstrated that in conditions of TORC1 inhibition, *ded1-mam* cells had increased translation of reporters with unstructured 5’ UTRs compared to wild-type *DED1*, and conversely, the mutants displayed decreased translation of structured reporters in the same conditions. Furthermore, these results may suggest that the rapamycin-resistant phenotype of the *ded1-mam* cells is at least partially due to increased translation of mRNAs with relatively unstructured 5’ UTRs that encode proteins that support growth. Such a model could also suggest how the *DDX3X/DED1* mutations affect oncogenesis in medulloblastoma.

To validate this model, we tested whether previously identified mRNA targets of Ded1 with relatively unstructured 5’ UTRs showed increased levels in *ded1-mam* mutants under stress (Supplementary Figure S4). Drawing from previously published ribosome sequencing, we arranged a list of mRNAs dependent on Ded1 in non-stress conditions by the structural complexity of their 5’ UTRs, as determined by PARS (Parallel Analysis of RNA Structure) score (21, 44). To then focus on genes linked to TOR inactivation, we retained only those that are annotated in the Saccharomyces Genome Database as displaying altered growth in rapamycin-treated cells when mutated. This left 102 genes that were arranged in ascending order by PARS score (Figure 5A). Taking the relatively unstructured half of these genes (PARS scores ≤ *PTC4*), we then focused on those that were annotated as rapamycin-sensitive upon deletion, implying that these genes contribute to growth and/or rapamycin resistance, and also had known functions related to stress response pathways. From the resulting list of 17, we selected 3 “pro-growth” genes to test for changes in the *ded1-mam* cells following rapamycin treatment: *CST6*, *VMA5*, and *EDE1*.

**Figure 5:**
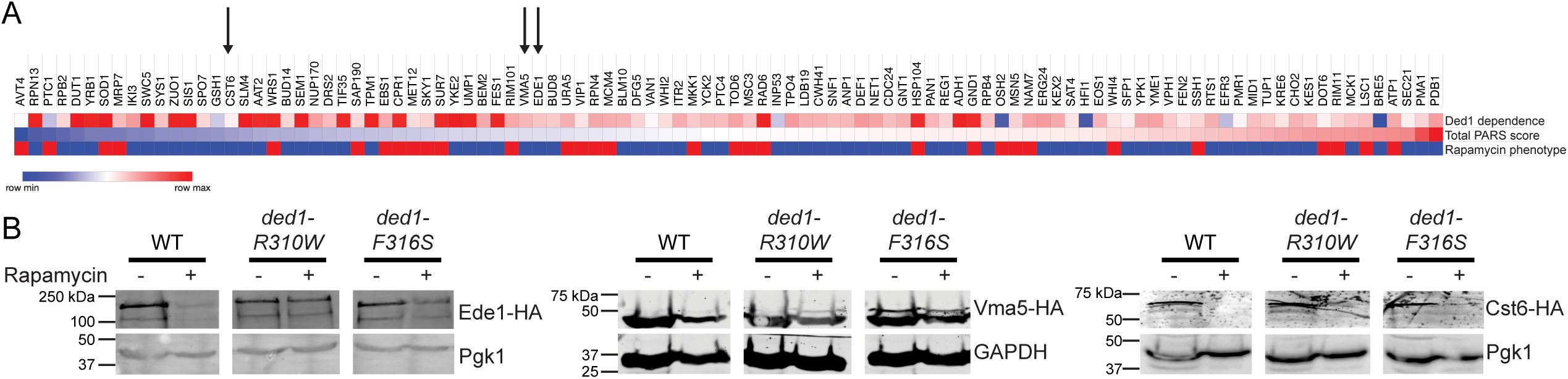
Medulloblastoma-associated mutants of DDX3X/DED1 have increased protein levels of known *DED1* targets with unstructured 5’ UTRs following TOR inactivation. (A) Heatmap showing the 102 Ded1-dependent genes that were annotated for rapamycin growth phenotypes, arranged in ascending order of their PARS score. Ded1 dependence is shown as the change in translation efficiency in a *ded1* temperature sensitive mutant where red is more sensitive to Ded1 perturbation. Total PARS score is shown where blue indicates less structure. Rapamycin-sensitive growth in a null mutant of the indicated gene is shown as blue, while rapamycin-resistance is shown in red. (B) Pro-growth genes with relatively unstructured 5’ UTRs identified to be rapamycin-resistant were tagged with 3xHA tag, then protein levels were examined in wild-type *DED1, ded1-R310W*, and *ded1-F316S*. Cells were treated with rapamycin (200 ng/ml), then protein extracts were run on SDS-PAGE and blotted using anti-HA with anti-Pgk1 and anti-GAPDH as loading controls. Representative images are shown.

To confirm our luciferase reporter results above, we integrated an HA tag into these putative Ded1 target genes in wild-type *DED1* cells and two *ded1-mam* strains (*ded1-R310W* and *-F316S*). After treating these cells with rapamycin for 4 hours, protein extracts were prepared and run on SDS-PAGE gels to analyze levels of the respective Ded1 targets in untreated and rapamycin-treated conditions. Consistent with our hypothesis, we observed an increase in the levels of each of the three proteins in the *ded1-mam* strains relative to wild-type *DED1* on rapamycin treatment (Figure 5B). This result implies that these pro-growth genes that have relatively unstructured 5’ UTRs may be translated at an increased rate following TOR inhibition, which would contribute to increased cell growth in the *ded1-mam* mutants relative to wild-type *DED1*. These observations support a model where, in medulloblastoma, mutated DDX3X upregulates the expression of unstructured, pro-growth genes, thus resulting in cancer progression and contributing to medulloblastoma pathology.

## Discussion

In this work, we have attempted to elucidate the molecular mechanism by which *DDX3X* mutations contribute to medulloblastoma progression. Using the genetically tractable yeast model, we introduced conserved medulloblastoma-associated mutations from mammalian *DDX3X* into the yeast homolog *DED1* (*ded1-mam*). In our prior studies with the *ded1-mam*, we discovered that they are not inactivating/null mutations and did not consistently reduce DDX3X/Ded1 function in different assays of activity. We observed significant variability in growth and translation defects in the mutants, suggesting that they are not simple, straightforward hypomorphic mutations. We found that the growth defects in *ded1-mam* were best correlated with defects in their 5’ UTR unwinding activity, i.e., these mutants had specific translation defects (31). However, this work focused on DDX3X/Ded1’s role under nutrient-rich, optimum conditions and did not take into account the role of DDX3X/Ded1 in stress responses.

In our current work, we expanded on this prior study and showed that roughly two-thirds of the mutants demonstrate rapamycin-resistant growth phenotypes (Figures 1B, S1, S2, and Table 1). Previously, we showed that many of the *ded1-mam* mutants had cold-sensitive growth defects in the absence of other stress, consistent with other studies and likely due to defects in helicase activity (31, 32, 45). Interestingly, here we observed a correlation between the two phenotypes (Figure S3 and Table 1). The mildly temperature-sensitive mutants (marked as (+++) in Table 1) were 44% rapamycin-resistant, the moderately temperature-sensitive mutants (++) were 75% rapamycin-resistant, and the extremely temperature-sensitive mutants (+) were 100% rapamycin-resistant (Figure S3). These results suggest that moderate defects in Ded1’s helicase activity might be behind this stress-resistance phenotype.

Numerous studies have established the critical role of DDX3X/Ded1 in translation (10, 13). Our prior work characterizing the *ded1-mam* mutants showed that while general, bulk translation was defective in some, this phenotype was highly inconsistent across a broad subset of the mutations (31). Here, we observed that relative to wild-type *DED1,* in the absence of rapamycin, the tested *ded1-mam* show decreased bulk translation, consistent with these previous results (Figure 2). In contrast, under rapamycin treatment, their translation profiles look similar to wild-type *DED1*. Thus, the fold-change values in *ded1-mam*’s translation profiles are reduced (Figure 2G). However, given the similarity in the P/M ratios in rapamycin, it seems unlikely that changes in bulk translation, whether increased or decreased, are responsible for the *ded1-mam* growth phenotypes. Rather, it is more likely that specific changes in translation contribute to the rapamycin-resistance phenotype.

Our previous work has also shown that under stress conditions, Ded1 triggers dissociation of eIF4G1 from mRNPs, resulting in the co-degradation of both factors (24). However, in a *ded1* mutant that deletes the eIF4G-interaction domain (*ded1-*Δ*CT*), this co-degradation is reduced, resulting in higher Ded1 and eIF4G1 levels and facilitating continued translation. Likewise, here we also observed higher levels of Ded1 and eIF4G1 in *ded1-mam* strains relative to wild-type *DED1* following TOR inhibition (Figure 3). Taken alone, this result suggests a mechanism similar to that observed in the *ded1-*Δ*CT* of decreased translation repression under stress. However, taken together with the polysome profiles in Figure 2, these results reinforce the implication that specific changes in translation are responsible for the rapamycin-resistance of *ded1-mam* and suggest that the levels of eIF4G1 and the mutant ded1 may also contribute to these differences (see Figure 6 below).

**Figure 6:**
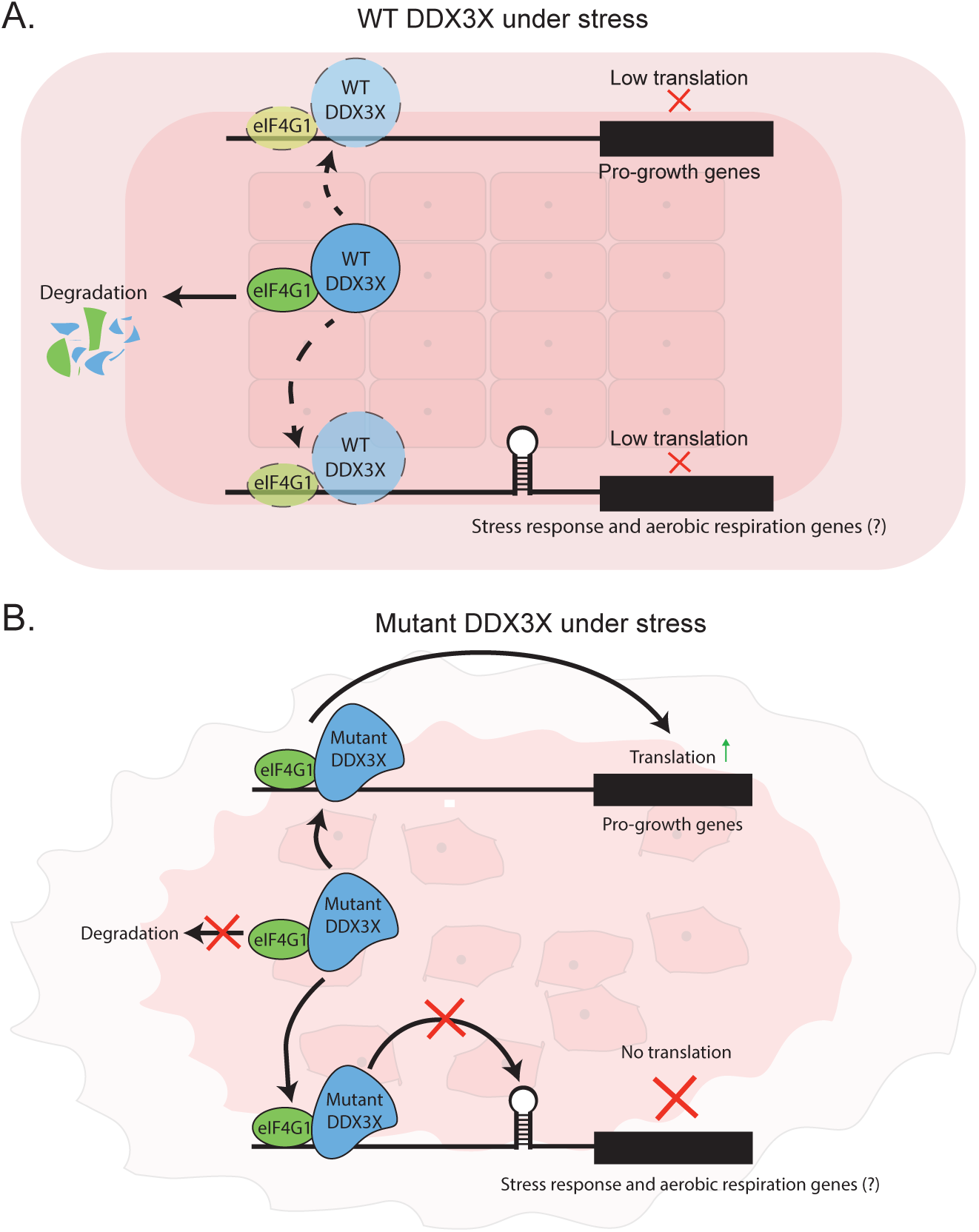
Model for the role of DDX3X/Ded1 in medulloblastoma progression. (A) Normally, when cells encounter stress conditions, wild-type DDX3X will cause eIF4G1 dissociation from translation complexes, and both eIF4G1 and DDX3X will subsequently degrade. The low levels of both proteins will downregulate translation for both pro-growth mRNAs with unstructured 5’ UTRs and mRNAs with structured 5’ UTRs, which may include inhibitors of the stress response and/or aerobic respiration genes. Some mRNAs with more complex regulatory mechanisms will presumably still be translated (not pictured). (B) However, in *ddx3x-mam* mutant cells, the aberrant DDX3X protein does not degrade eIF4G1, allowing continued translation of unstructured, pro-growth genes and helping tumor cells bypass stress-induced growth inhibition. Conversely, the mutant DDX3X is not able to support translation of mRNAs with structured 5’ UTRs, possibly resulting in a partial stress response and changes in energy metabolism in the tumor.

Prior research has also shown that DDX3X/Ded1’s function in unwinding secondary structures in the 5’ UTR for facilitating start site scanning is affected by these medulloblastoma mutations (31). Therefore, we hypothesized that this deficiency contributes to the rapamycin-resistant phenotypes in the *ded1-mam* cells. As shown in Figure 4, *ded1-mam* mutants translated the structured 5’ UTR reporters at a decreased rate relative to wild-type, regardless of the presence of rapamycin. This result extends our prior study, suggesting that the *ded1-mam* have difficulty unwinding structures in 5’ UTRs under any conditions. However, for the unstructured 5’ UTR reporters, we saw that they were translated in *ded1-mam* at a rate higher than wild-type *DED1* when treated with rapamycin. These observations suggest that the rapamycin-resistant growth of the *ded1-mam* cells is due to the increased translation of mRNAs with relatively unstructured 5’ UTRs that promote growth and would normally be repressed under stress conditions. Conversely, mRNAs with structured 5’ UTRs that show decreased translation may be stress-response or growth-inhibitory genes.

As discussed above, our data suggest that the *DDX3X/DED1* medulloblastoma-associated mutations affect the translatome in a complex manner, probably via the effects of DDX3X’s ability to unwind 5’ UTR structures. This model is also consistent with that of Oh et al., who likewise proposed that there would be specific subsets of genes that would be selectively translated during stress conditions in the medulloblastoma-associated *DDX3X-R543H* mutant (33). To test whether our luciferase reporter findings can further inform this model, we identified genes related to growth and stress responses from previously identified Ded1-targeted mRNAs (37). Following TOR inhibition, we observed that the protein levels of pro-growth genes that had relatively unstructured 5’ UTRs showed significant upregulation in *ded1-mam* relative to wild-type cells, implying that these mRNAs have increased translation in the mutants compared to wild-type cells in these conditions (Figure 5).

Taken all together, our results suggest a model wherein Ded1/DDX3X normally promotes translation repression of pro-growth genes during stress conditions, at least partially through dissociation and degradation of eIF4G1, in order to support the redirection of resources from cell growth to cell survival (Figure 6, top). However, medulloblastoma-associated mutants of Ded1 are not able to repress translation, and eIF4G1 is maintained at higher levels, promoting continued growth even during rapamycin treatment (Figure 6, bottom). Extrapolating to cancer, mutation of *DDX3X* would likewise promote the translation of pro-growth mRNAs (with unstructured 5’ UTRs), even in the interior of a tumor where cancer cells have limited nutrients and other stress conditions are present (46). Correspondingly, other subsets of mRNAs, such as inhibitors of the stress response and/or enzymes involved in aerobic respiration, that have structured 5’UTRs will be further downregulated in cells with mutant Ded1/DDX3X due to the defects in its helicase activity. In medulloblastoma, this would result in the upregulation of stress-response genes (antioxidants, chaperones, etc.) and/or metabolic changes in the tumor, respectively. Given the identified roles of DDX3X in neuron generation and differentiation (28–30), mRNAs involved in these processes may be regulated similarly in a *DDX3X* mutant to promote medulloblastoma progression.

This model suggests that *DDX3X/DED1* does not fit neatly into the categories of tumor suppressor or proto-oncogene, displaying properties of both depending on the target mRNA. These selective effects, while broadly consistent with other studies of the medulloblastoma mutants, have not been widely demonstrated (31, 33). As mentioned above, Oh et al. proposed selective effects during stress conditions, specifically oxidative stress, although they observed only moderate defects in one mutant (33). Another study used a conditional knockout model of *DDX3X* to suggest that it acts as a tumor suppressor by moderating the effects of upregulated Wnt signaling, which include cell death, cell cycle arrest, and “oncogenic stress” (30). However, a *ddx3x* knockout may not fully recapitulate the effects of what appear to be complex hypomorphic mutations in medulloblastoma. Furthermore, only RNA levels were examined rather than translation itself, so some caution is needed in interpreting these results. Our data extends these results by both suggesting a potential mechanism and by proposing an additional layer of regulation. Along these lines, Valentin-Vega et al., as well as others, observed that some of the medulloblastoma mutants exhibited increased stress granule (SG) formation (31, 33, 34). Although increased SGs are linked to cancer, their role remains somewhat unclear (47). In this study, we note that rapamycin does not induce SG formation in yeast, so the *ded1-mam* phenotypes we have observed presumably are not due to SGs.

This study defines the role of *ded1-mam* mutants in mediating the cellular stress response and provides new mechanistic insight into the conserved function of DDX3X/Ded1 in cancer. Our findings suggest that DDX3X/Ded1 selectively unwinds and promotes the translation of specific mRNAs based on 5′UTR structure, a process modulated by eIF4G1 availability under stress conditions. Dysregulation of this mechanism in *DDX3X/DED1* mutants enables medulloblastoma cells to sustain translation and proliferate under adverse conditions, thereby contributing to tumor progression. Given these striking phenotypes in yeast cells, future studies should include validating these stress-adaptive phenotypes in human cancer cells, possibly by using CRISPR/Cas9 to generate the appropriate *ddx3x* mutants in neurons or neuronally-derived cell lines. Additionally, ribosome profiling of these mutants and verifications of *DDX3X*’s translational targets in such a model would be critical to furthering our knowledge of DDX3X’s precise role in tumor progression. Such insights could help us identify therapeutic targets for medulloblastoma and inform more improved strategies for targeted treatment.

### Experimental procedures

#### Strain and plasmid construction

Yeast strains and plasmids used are listed in Tables S1 and S2. The *ded1-mam* strains were generated as described previously (31). Ded1 target strains (TBY258, 262, 266) were generated by integrating a C-terminal 3x-HA tag marked with *HIS3* at the respective locus (*CST6, VMA5*, *EDE1*) into the yeast strain SWY4093 (*ded1::KAN +pDED1/URA3/CEN*). The *DED1* and *ded1-*mam plasmids were then shuffled into these parental strains. Yeast growth assays were performed by serial dilution as previously described on rich media (YPD) in the presence or absence of 200 ng/mL rapamycin (24).

#### Western blotting

Blotting for protein levels of Ded, eIF4G1 (Tif4631), and Ded1 targets was performed as previously described (36). Cells were grown to early-log phase and then subjected to rapamycin treatment (200 ng/ml) for 4 hours. Crude cell lysates were prepared by lysing the cells in 1.85M NaOH and 7.4% beta-mercaptoethanol, followed by trichloroacetic acid precipitation. Proteins were separated by SDS-PAGE and probed with antibodies specific against Ded1 (48), eIF4G1 (24), and anti-HA (12CA5) for Vma5, Ede1 and CST6 (Roche). Antibodies against Pgk1 and GAPDH (Invitrogen) were used as loading controls. Fluorescent secondary antibodies (IRDye 680RD Goat Anti-Rabbit IgG and IRDye 800CW Goat anti-Mouse IgG) were used to visualize bands on an Odyssey M imager (LI-COR Biosystems). For densitometry analysis, protein band intensity was measured with ImageJ/Fiji software. Ded1 and eIF4G1 were normalized to Pgk1 band intensity of the same sample. Statistical significance was calculated using Student’s t-test.

#### Polyribosome profiles

Polysome profiles were generated essentially as described previously (36). Briefly, wild-type *DED1, ded1-R310W,* and *ded1-F316S* were grown to mid-log phase, and 200 ng/ml of rapamycin was added for 40 minutes. Ribosomes were stalled on mRNAs via the addition of cycloheximide, and then cells were lysed using a bead beater (2x for 2.5 min). Lysates were loaded on a 7 to 47% sucrose density gradient and centrifuged for 2.5 hours at 38,000 rpm. RNA concentration of the samples was then monitored via absorbance at 254 nm during continuous fractionation. Polysome/monosome ratios were calculated by comparing the sum of the areas under the polysome peaks to the area under the 80S monosome peak in ImageJ/Fiji. Statistical significance was determined via Mann-Whitney non-parametric tests in GraphPad Prism.

#### Translation scanning assays

Scanning assays to study the translation of mRNAs with different 5’ UTR sequences were carried out essentially as described (37). Wild-type *DED1* and the indicated *ded1-mam* mutant cells were transformed with an unstructured *RPL41A* 5’ UTR-firefly luciferase reporter (pFJZ342) or a distal-stem-loop-containing version (ΔG = -3.7 kcal/mol, pFJZ623). Transformants were cultured in triplicate in early log phase and then treated with rapamycin (200 ng/ml) for 4 hours. Cell lysates were generated, and luciferase assays were performed with standard luciferin reagent (Promega) on a Glomax 20/20 luminometer (Promega) as previously described (31). Triplicates were normalized to their respective cell concentrations and averaged together for each biological replicate. Statistical significance was determined using Student’s unpaired t-test.

#### Selection of Ded1-dependent mRNAs

To identify unstructured mRNAs that are dependent on Ded1 activity, previously published ribosome sequencing data was used, and PARS (Parallel Analysis of RNA Structure) scores, as determined by Kertesz et al., were used as a proxy for structural complexity of the 5’UTR (37, 44). From a master list of 814 genes that were Ded1-dependent, 457 of them had designated PARS scores (Supplementary Figure S4). To narrow down this list further to select for genes involved in TOR inactivation, the Alliance Mine tool on the Saccharomyces Genome Database (SGD) was used to select for genes with a growth phenotype change in the presence of sirolimus/rapamycin. These filters gave 102 (out of the 457) genes annotated for increased or decreased growth in rapamycin-treated cells when mutated. These 102 genes were arranged in ascending order of their PARS score (see Figure 5A). The first half of these genes (PARS scores ≤ *PTC4*) were designated as relatively unstructured, and then genes for which ΔTE (translational efficiency) is negative were selected, narrowing the list to 32 genes. Genes wherein the null mutant was rapamycin-sensitive (the expected phenotype, given that increased translation of these genes in *ded1-mam* was predicted) left 19 genes. From this list, 3 stress-related and “pro-growth” genes (based on their phenotypes and known functions), *CST6*, *VMA5*, and *EDE1*, were selected for further analysis.

## Data availability

All data are contained within the article or supplementary information. Original data will be shared upon request of the corresponding author.

## Acknowledgments

The authors would like to thank Alan Hinnebusch for plasmids and Christopher West and Natalie Krahn for laboratory equipment and reagents. We would also like to thank David Garfinkel and members of the Bolger lab, particularly Allison Michael, for helpful discussions and feedback.

## Funding and additional information

This work was supported by a grant from the National Institute of General Medical Sciences (1R01-GM136827). The content is solely the responsibility of the authors and does not necessarily represent the official views of the National Institutes of Health or other funding organizations.

## Abbreviations

TOR: Target of Rapamycin
TORC1: Target of Rapamycin Complex 1
RNP: Ribonuceloprotein
eIF: eukaryotic initiation factor
P/M: polysome to monosome
PIC: preinitiation complex
SG: stress granule
UTR: untranslated region.

