## Supporting Information for "Medulloblastoma-associated mutations in the RNA helicase *DDX3X/DED1* cause defects in the translational response to TORC1 inhibition"

1. Supplementary table S1: Yeast strains used in this study
2. Supplementary table S2: Plasmids used in this study
3. Figure S1: Growth and rapamycin-sensitivity of DDX3X/DED1 medulloblastoma-associated mutations at 25°C.
4. Figure S2: Growth and rapamycin-sensitivity of DDX3X/DED1 medulloblastoma-associated mutations at 30°C.
5. Figure S3: Correlation between growth phenotypes and rapamycin-resistance of *ded1-mam* cells.
6. Figure S4: Selection of Ded1-dependent genes with relatively unstructured 5' UTRs.

**Supplementary table S1: Yeast strains used in this study**

| Strain name | Genotype | Source |
|---|---|---|
| W303 | *MATα ade2-1 ura3-1 his3-11,15 leu2-3,112 trp1-1 can1-100* | (49) |
| SWY4093 | *MATα ded1::KANr ade2-1 ura3-1 his3-11,15 leu2-3,112 trp1-1 can1-100 +pCEN/URA3/DED1* | (48) |
| TBY52 | *MATα ded1::KANr ade2-1 ura3-1 his3-11,15 leu2-3,112 trp1-1 can1-100 +pCEN/LEU2/DED1* (pSW3619) | (24) |
| TBY 121 | *MATα ded1::KANr ade2-1 ura3-1 his3-11,15 leu2-3,112 trp1-1 can1-100 +pCEN/LEU2/ded1-ΔCT* | (24) |
| SWY4277 | *MATα ded1::KANr ade2-1 ura3-1 his3-11,15 leu2-3,112 trp1-1 can1-100 +pCEN/LEU2/ded1-120* | (48) |
| TBY 258 | *MATα ded1::KANr CST6:3XHA:HIS3 ade2-1 ura3-1 his3-11,15 leu2-3,112 trp1-1 can1-100 +pCEN/URA3/DED1* | This study |
| TBY 259 | *MATα ded1::KANr CST6:3XHA:HIS3 ade2-1 ura3-1 his3-11,15 leu2-3,112 trp1-1 can1-100 +pCEN/LEU2/DED1* | This study |
| TBY 260 | *MATα ded1::KANr CST6:3XHA:HIS3 ade2-1 ura3-1 his3-11,15 leu2-3,112 trp1-1 can1-100 +pCEN/LEU2/ded1-R310W* | This study |
| TBY 261 | *MATα ded1::KANr CST6:3XHA:HIS3 ade2-1 ura3-1 his3-11,15 leu2-3,112 trp1-1 can1-100 +pCEN/LEU2/ded1-F316S* | This study |
| TBY 262 | *MATα ded1::KANr VMA5:3XHA:HIS3 ade2-1 ura3-1 his3-11,15 leu2-3,112 trp1-1 can1-100 +pCEN/URA3/DED1* | This study |
| TBY 263 | *MATα ded1::KANr VMA5:3XHA:HIS3 ade2-1 ura3-1 his3-11,15 leu2-3,112 trp1-1 can1-100 +pCEN/LEU2/DED1* | This study |
| TBY 264 | *MATα ded1::KANr VMA5:3XHA:HIS3 ade2-1 ura3-1 his3-11,15 leu2-3,112 trp1-1 can1-100 +pCEN/LEU2/ded1-R310W* | This study |
| TBY 265 | *MATα ded1::KANr VMA5:3XHA:HIS3 ade2-1 ura3-1 his3-11,15 leu2-3,112 trp1-1 can1-100 +pCEN/LEU2/ded1-F316S* | This study |
| TBY 266 | *MATα ded1::KANr EDE1:3XHA:HIS3 ade2-1 ura3-1 his3-11,15 leu2-3,112 trp1-1 can1-100 +pCEN/URA3/DED1* | This study |
| TBY 267 | *MATα ded1::KANr EDE1:3XHA:HIS3 ade2-1 ura3-1 his3-11,15 leu2-3,112 trp1-1 can1-100 +pCEN/LEU2/DED1* | This study |
| TBY 268 | *MATα ded1::KANr EDE1:3XHA:HIS3 ade2-1 ura3-1 his3-11,15 leu2-3,112 trp1-1 can1-100 +pCEN/LEU2/ded1-R310W* | This study |
| TBY 269 | *MATα ded1::KANr EDE1:3XHA:HIS3 ade2-1 ura3-1 his3-11,15 leu2-3,112 trp1-1 can1-100 +pCEN/LEU2/ded1-F316S* | This study |

*ded1-mam* list (*MATα ded1::KANr ade2-1 ura3-1 his3-11,15 leu2-3,112 trp1-1 can1-100 +pCEN/LEU2/ded1-mam*):

| **Strain name** | **Genotype** | **Source** |
|---|---|---|
| TBY53 | *-R285C(pTB38)* | (31) |
| TBY54 | *-F316S (pTB39)* | (31) |
| TBY55 | *-P526L (pTB42)* | (31) |
| TBY56 | *-R492H (pTB43)* | (31) |
| TBY57 | *-T234M (pTB40)* | (31) |
| TBY58 | *-R335C (pTB41)* | (31) |
| TBY59 | *-S371F (pTB44)* | (31) |
| TBY60 | *-A184P (pTB45)* | (31) |
| TBY61 | *-R235K (pTB59)* | (31) |
| TBY62 | *-D313H (pTB52)* | (31) |
| TBY63 | *-M329R (pTB54)* | (31) |
| TBY64 | *-L312F (pTB48)* | (31) |
| TBY65 | *-D288V (pTB49)* | (31) |
| TBY66 | *-G488V (pTB50)* | (31) |
| TBY67 | *-R433C (pTB51)* | (31) |
| TBY68 | *-T166A (pTB55)* | (31) |
| TBY69 | *-V168L (pTB56)* | (31) |
| TBY70 | *-R486H (pTB67)* | (31) |
| TBY71 | *-L393D (pTB66)* | (31) |
| TBY72 | *-G189V (pTB57)* | (31) |
| TBY73 | *-T193A (pTB63)* | (31) |
| TBY74 | *-G262V (pTB60)* | (31) |
| TBY75 | *-L286V (pTB62)* | (31) |
| TBY76 | *-M339I (pTB71)* | (31) |
| TBY77 | *-V364L (pTB74)* | (31) |
| TBY78 | *-T490M (pTB68)* | (31) |
| TBY79 | *-R310W (pTB73)* | (31) |
| TBY80 | *-G494R (pTB65)* | (31) |
| TBY81 | *-E236Q (pTB64)* | (31) |
| TBY82 | *-A352S (pTB76)* | (31) |
| TBY83 | *-V454M (PTB78)* | (31) |
| TBY87 | *-G462V (pTB88)* | (31) |
| TBY88 | *-D326V (pTB89)* | (31) |

**Supplementary table S2: Plasmids used in this study**

| Plasmid Name | Description | Source |
|---|---|---|
| pRS315 | *CEN/LEU2* | (50) |
| pSW3619 | *CEN/LEU2/DED1* | (48) |
| pTB38 | *CEN/LEU2/ded1-R285C* | (31) |
| pTB39 | *CEN/LEU2/ded1-F316S* | (31) |
| pTB40 | *CEN/LEU2/ded1-T234M* | (31) |
| pTB41 | *CEN/LEU2/ded1-R335C* | (31) |
| pTB42 | *CEN/LEU2/ded1-P526L* | (31) |
| pTB43 | *CEN/LEU2/ded1-D464Y* | (31) |
| pTB44 | *CEN/LEU2/ded1-S371F* | (31) |
| pTB45 | *CEN/LEU2/ded1-A184P* | (31) |
| pTB46 | *CEN/LEU2/ded1-R492H* | (31) |
| pTB47 | *CEN/LEU2/ded1-G284E* | (31) |
| pTB48 | *CEN/LEU2/ded1-L312F* | (31) |
| pTB49 | *CEN/LEU2/ded1-D288V* | (31) |
| pTB50 | *CEN/LEU2/ded1-G488A* | (31) |
| pTB51 | *CEN/LEU2/ded1-R433C* | (31) |
| pTB52 | *CEN/LEU2/ded1-D313H* | (31) |
| pTB53 | *CEN/LEU2/ded1-G261V* | (31) |
| pTB54 | *CEN/LEU2/ded1-M329R* | (31) |
| pTB55 | *CEN/LEU2/ded1-T166A* | (31) |
| pTB56 | *CEN/LEU2/ded1-V168L* | (31) |
| pTB57 | *CEN/LEU2/ded1-G189V* | (31) |
| pTB59 | *CEN/LEU2/ded1-R235K* | (31) |
| pTB60 | *CEN/LEU2/ded1-G262V* | (31) |
| pTB61 | *CEN/LEU2/ded1-H485Y* | (31) |
| pTB62 | *CEN/LEU2/ded1-L286V* | (31) |
| pTB63 | *CEN/LEU2/ded1-T193A* | (31) |
| pTB64 | *CEN/LEU2/ded1-E236Q* | (31) |
| pTB65 | *CEN/LEU2/ded1-G494R* | (31) |
| pTB66 | *CEN/LEU2/ded1-L393!1* | (31) |
| pTB67 | *CEN/LEU2/ded1-R486H* | (31) |
| pTB68 | *CEN/LEU2/ded1-T490M* | (31) |
| pTB70 | *CEN/LEU2/ded1-I374!1* | (31) |
| pTB71 | *CEN/LEU2/ded1-M339I* | (31) |
| pTB72 | *CEN/LEU2/ded1-T343P* | (31) |
| pTB73 | *CEN/LEU2/ded1-R310W* | (31) |

| | | |
|---|---|---|
| pTB74 | *CEN/LEU2/ded1-V364L* | (31) |
| pTB76 | *CEN/LEU2/ded1-A352S* | (31) |
| pTB77 | *CEN/LEU2/ded1-A4591* | (31) |
| pTB78 | *CEN/LEU2/ded1-V454M* | (31) |
| pTB88 | *CEN/LEU2/ded1-G462V* | (31) |
| pTB89 | *CEN/LEU2/ded1-D326V* | (31) |
| pTB90 | *CEN/LEU2/ded1-Q240H* | (31) |
| pTB109 | *CEN/LEU2/ded1-V176S* | (31) |
| pTB136 | *CEN/LEU2/ded1-ΔCT* | (24) |
| *ded1-120* | *CEN/LEU2/ded1-120* | (13) |
| pFJZ342 | *CEN/URA3/5’UTR-RPL41A+CAA(23)-LUC* | (21) |
| pFJZ623 | *CEN/URA3/5’UTR-RPL41A+CAA(23)+distal-stem-loop(3.7)-LUC* | (21) |

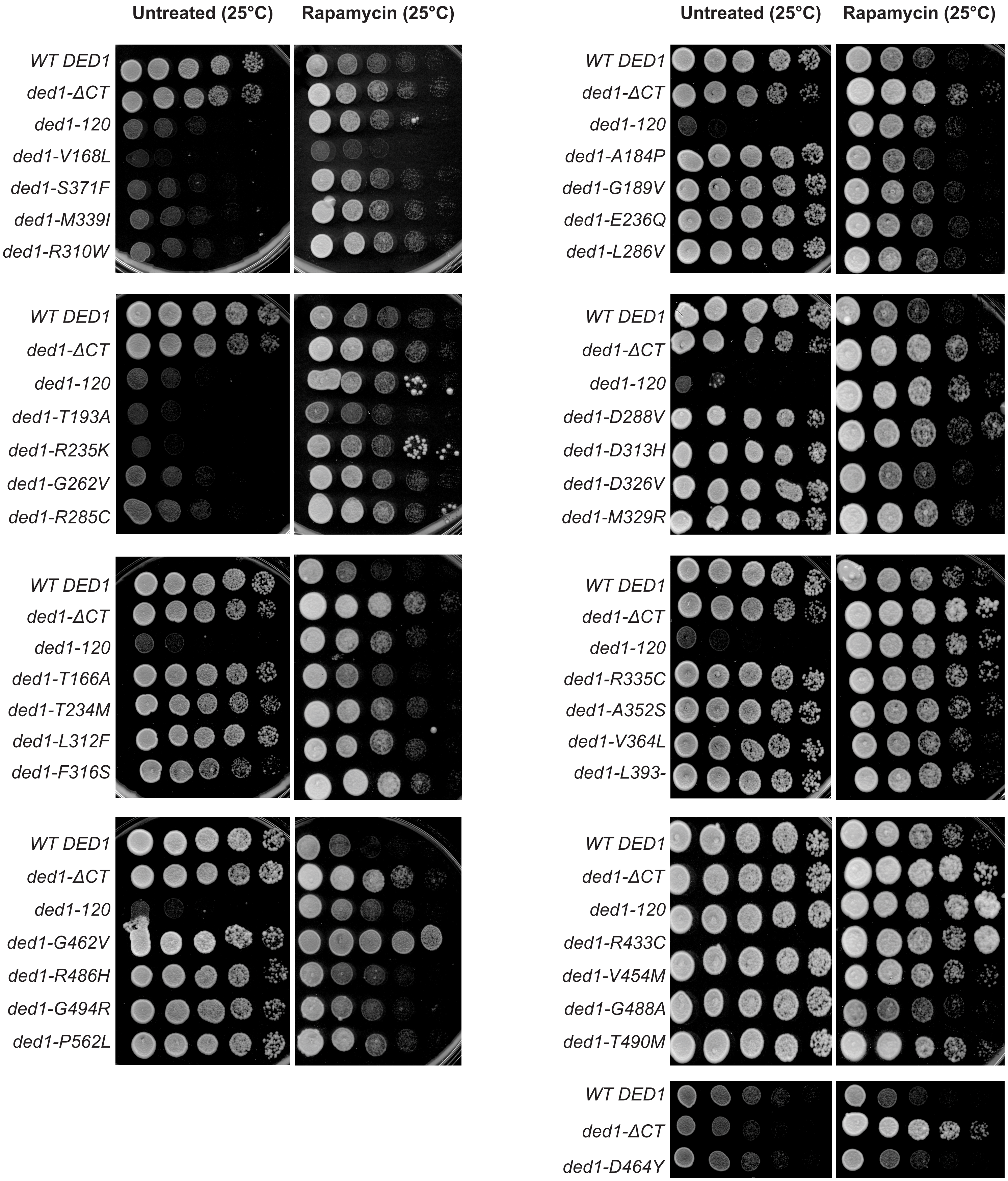


**Figure S1: Growth and rapamycin-sensitivity of *DDX3X/DED1* medulloblastoma-associated mutations at 25°C.** Wild-type *DED1* cells and the indicated *ded1-mam* mutants were serially diluted onto nutrient-rich agar in the presence or absence of rapamycin (200 ng/ml) and incubated at 25°C.

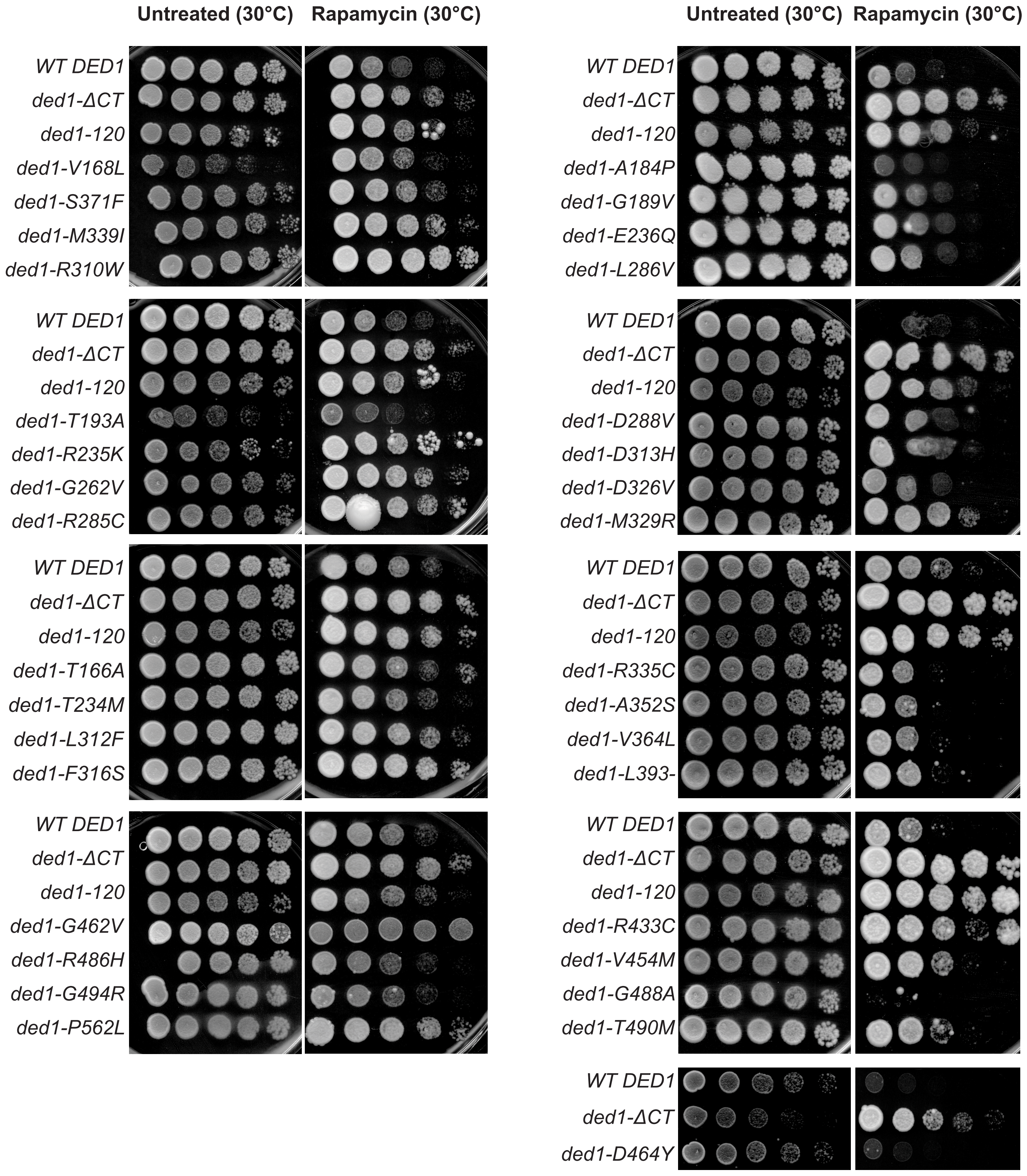


**Figure S2: Growth and rapamycin-sensitivity of *DDX3X/DED1* medulloblastoma-associated mutations at 30°C.** Wild-type *DED1* cells and the indicated *ded1-mam* mutants were serially diluted onto nutrient-rich agar in the presence or absence of rapamycin (200 ng/ml) and incubated at 30°C.

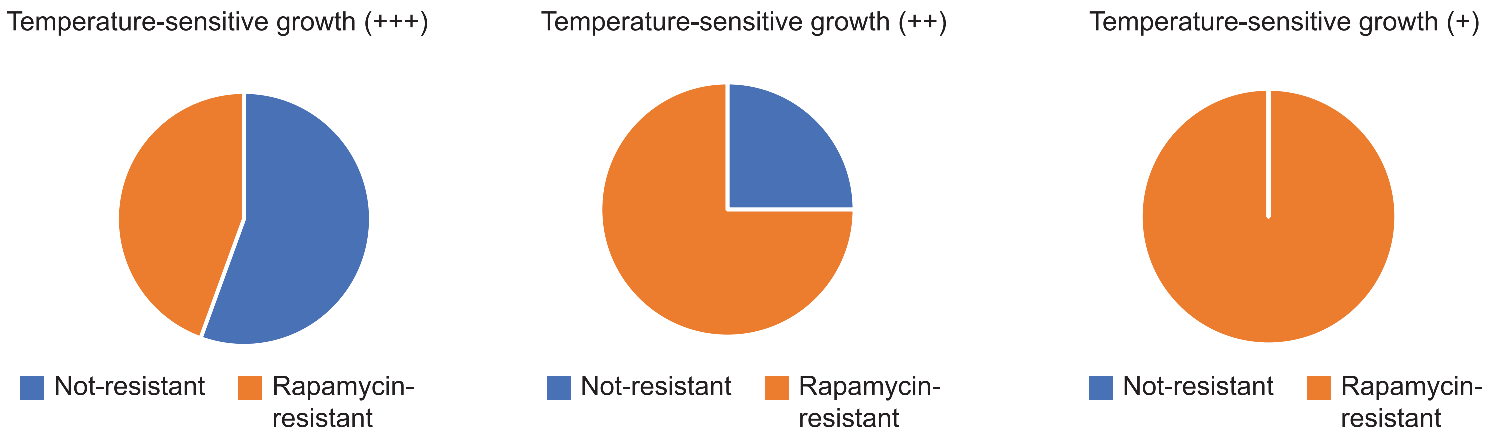


**Figure S3: Correlation between growth phenotypes and rapamycin-resistance of *ded1-mam* cells.** All non-lethal ded1-mam mutants were sorted into 3 categories based on their temperature sensitivity on rich media: normal growth at 25°C (+++), moderate growth (++), and poor growth (+). The percentage of mutants showing rapamycin-resistant growth in each category was then graphed. (+++) 8 out of 18 were rapamycin-resistant; (++) 6 of 8; for (+) 7 of 7.

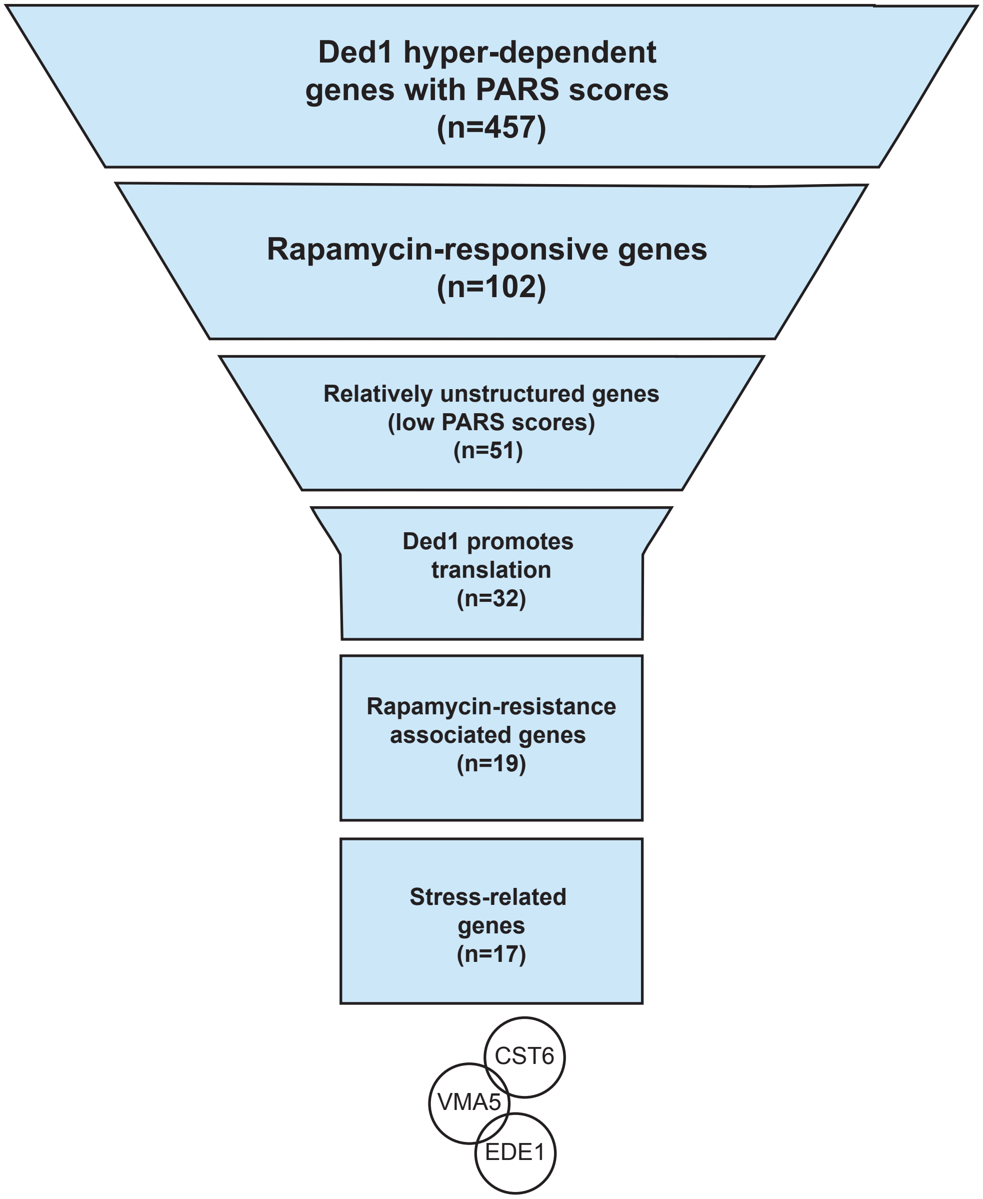


**Figure S4: Selection of Ded1-dependent genes with relatively unstructured 5' UTRs.** From a list of 814 Ded1-dependent genes, we selected 3 highly Ded1-dependent gene with relatively unstructured 5' UTRs that are rapamycin-resistance associated and stress-related.
